# RPG: A low-cost, open-source, high-performance solution for displaying visual stimuli

**DOI:** 10.1101/2020.03.05.979724

**Authors:** Vivian Imbriotis, Adam Ranson, William M Connelly

## Abstract

The development of new high throughput approaches for neuroscience such as high-density silicon probes and 2-photon imaging have led to a renaissance in visual neuroscience. However, generating the stimuli needed to evoke activity in the visual system still represents a non-negligible difficulty for experimentalists. While several widely used software toolkits exist to deliver such stimuli, they all suffer from some shortcomings. Primarily, the hardware needed to effectively display such stimuli comes at a significant financial cost, and secondly, triggering and/or timing the stimuli such that it can be accurately synchronized with other devices requires the use of legacy hardware, further hardware, or bespoke solutions.

Here we present RPG, a Python package written for the Raspberry Pi, which overcomes these issues. Specifically, the Raspberry Pi is a low-cost, credit card sized computer with general purpose input/output pins, allowing RPG to be triggered to deliver stimuli and to provide real-time feedback on stimulus timing. RPG delivers stimuli at >60 frames per second and the feedback of frame timings is accurate to 10s of microseconds.

We provide a simple to use Python interface that is capable of generating drifting sine wave gratings, Gabor patches and displaying raw images/video.

## 1. Introduction

Using a computer to control the presentation of stimuli is a common task in visual neuroscience. As such, several software packages have been developed to deliver stimuli, most notably Psychophysics Toolbox (Brainard, 1997) and PsychoPy (Peirce, 2007). These packages can provide a diverse range of stimuli and are essentially capable of running entire experiments directly. However, they do have two significant limitations. Firstly, they are recommended to be run on relatively costly modern computers with graphics cards. Secondly, triggering the software from other sources or getting the software to trigger other equipment relies on a collection of non-ideal solutions. A primary option is the use of parallel or serial ports (e.g. Nagel & Spüler 2018), which are both legacy hardware, being largely supplanted by USB. A second option is to use external data acquisition hardware (e.g. Schneider et al.,2018), however, this further complicates the overall code, and increases costs. Finally, bespoke solutions such as using photodiodes to monitor screens can be used to collect frame timing (e.g. Aller et al., 2015). The diversity of computing hardware further complicates matters, such that code which utilizing this software may run perfectly on one computer, but result in missed timings or dropped frames on another.

Here we present RPG, a high performance, low cost, open source solution for displaying drifting gratings, Gabor patches and raw images/video. RPG is written in a combination of the languages Python and C, with Python providing a user-friend way to interface with the underlying code written in C for performance. RPG runs exclusively on the Raspberry Pi, a single-board computer, running the Raspbian Linux distribution. The Raspberry Pi features general purpose input/output (GPIO) pins, which are used by RPG digital input/outputs, specifically to trigger the display of stimuli and to provide output for the exact timing of frames. By running on standardized, low-cost hardware that comes equipped with GPIOs, we believe that some scientists will find RPG to be a superior option for displaying visual stimuli.

RPG is not intended as a replacement for other psychophysics toolboxes, as it lacks many of the functions that these excellent packages provide. However, for scientists who simply need to display drifting/stationary gratings, Gabor patches or raw video/images, we believe RPG will provide a simpler, more robust, and more reproducible solution than existing software.

## 2. Methods

### 2.1 Raspberry Pi

RPG has been developed and tested on the Model B Raspberry Pi 3 and 4, which are powered by a quad core Cortex A53 (1.2 GHz) and A72 (1.5 GHz) with 1GB and 4GB of RAM and running the default Raspbian GNU/Linux 9 “Stretch” and Raspian GNU/Linux 10 “Buster” operating systems respectively. The monitor was a Dell U2415.

### 2.2 Windows PC

Bench-marking of packages with similar functionality was performed on an Intel Core i7-8700k running at 3.7 GHz with 32GB Ram and a NVIDA GeForce 1080 Ti. Psychtoolbox (v3.0.15) was run in Matlab (R2019a) and PsychoPy 3.2.0 was run in Python 3.7.1. The same computer was used to monitor RPG performance, with a PCIe-6321 data acquisition device, (National Instruments) sampling at 80 kHz.

### 2.3 Animal surgery

All experimental procedures were carried out in accordance with the UK Animals (Scientific Procedures) Act 1986 and European Commission directive 2010/63/EU or with guidelines approved by the Animal Ethics Committee of the University of Tasmania.

All surgical procedures were conducted under aseptic conditions. For imaging experiments, prior to cranial window surgery, animals were administered with the antibiotic Baytril (5mg/kg, s.c.) and the anti-inflammatory drugs Rimadyl (5mg/kg, s.c.) and Dexamethasone (0.15mg/Kg, i.m.). Anesthesia was induced at a concentration of 4% Isoflurane in oxygen, and maintained at 1.5-2% for the duration of the surgery. Once anesthetised, animals were secured in a stereotaxic frame (David Kopf Instruments, Tujunga, CA, USA) and the scalp and periosteum were removed from the dorsal surface of the skull. A head plate was attached to the cranium using dental cement (Super Bond, C&B), with an aperture approximately centred over the right V1. A 3mm circular craniotomy was then made, centred −3.4 mm posterior and 2.8 mm lateral from bregma of the right hemisphere. After craniotomy a virus was injected into V1 (depth = 250µm, 40nl at 3 sites) to drive expression of GCaMP6F (AAV1.Syn.GCaMP6f.WPRE.SV40; titre after dilution 2×10^11^ GC/ml). The craniotomy was closed with a glass insert made from 3 layers of circular glass (#1 thickness; 1×5 mm, 2×3 mm diameter) bonded together with optical adhesive (Norland Products; catalogue no. 7106). The window was then sealed with dental cement. After surgery, all animals were allowed at least 1 week to recover before being imaged.

In vivo 2-photon imaging was performed using a resonant scanning microscope (Thorlabs, B-Scope) with a 16x 0.8NA objective (Nikon). GCaMP6 was excited at 980nm using a Ti:sapphire laser (Coherent, Chameleon) with a maximum laser power at sample of 50mW. Data were acquired at approximately 60Hz and averaged, resulting in a framerate of approximately 10Hz. For all imaging experiments animals were head-fixed and free to run on a cylindrical treadmill. Imaging was acquired using custom DAQ code (Matlab) and a DAQ card (NI PCIe-6323, National Instruments).

For electrophysiology, a 6 week old, male, C57BL/6j mouse was anaesthetized with Urethane (1 mg/kg) and chlorprothixene (5 mg/kg) in 0.9% saline I.P and maintained at 36-37 °C. A 3mm craniotomy was performed over the right visual cortex, which was half covered with a hemisected 5 mm coverslip adhered to the skull with dental cement. Under 2-photon guidance, a pipette was lowered into layer 2/3 and Cal520-AM (1 mM) and Sulforhodamine 101 (50 µM) was pressure ejected to label neurons and astrocytes respectively. After at least 30 minutes, a 5 MΩ electrode coated in quantum dots (Andrásfalvy et al., 2014) containing (in mM) KMeSo4 (120) KCl (10), HEPES (10), Mg-ATP (4), Na_2_-GTP (0.3), Na2-phosphocreatine (10) (pH 7.25–7.30 pH; 290–300 mOsm) was lowered into the same region and fluorescent neurons were targeted for whole cell recording. Recordings were made with AxoClamp 700B (Molecular Devices), filtered a 5 kHz, digitized at 50 kHz with an ITC-18 interface under the control of AxoGraph (AxoGraph).

## 3. Results

### 3.1 What RPG provides

RPG, along with the Raspberry Pi hardware, provides a convenient platform to generate and display visual stimuli (Fig 1). RPG is available to download at https://github.com/bill-connelly/rpg, along with a full code reference. RPG allows users to generate drifting or stationary gratings and Gabor patches and save them to disk. Also, arbitrary images and videos can be converted from a raw format, to an RPG format. These gratings, videos or images can then be loaded to memory where their display can be triggered programmatically or by a 3.3 V signal applied to a user selectable pin on the general purpose input/output (GPIO) header. The timings of the generated frames can be monitored by sampling the output of GPIO on physical pin 12, which outputs a 2ms long, 3.3 V pulse. The delivery of this pulse is tied to a successful write of the image to the framebuffer and the v-sync of the monitor.

**Figure 1.**
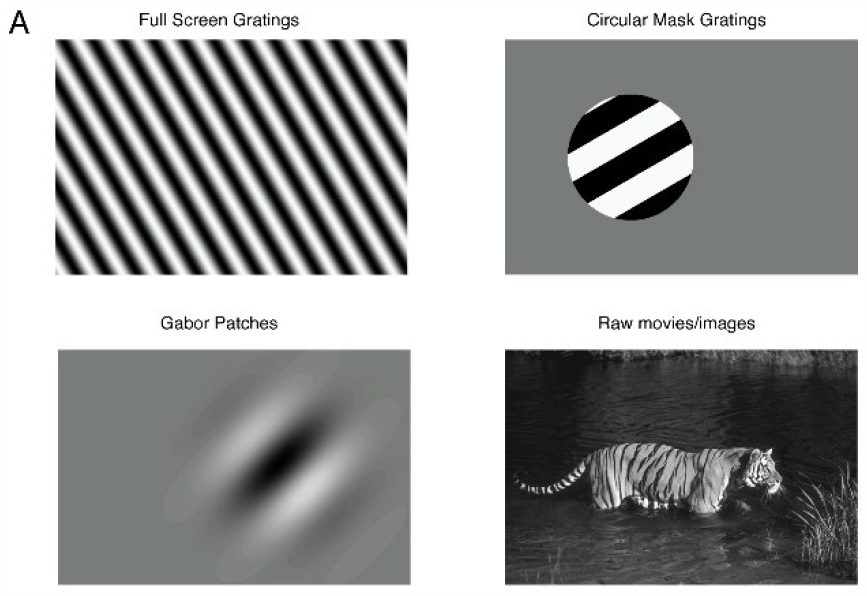
RPG can generate and display full screen or circular masked drifting or stationary gratings made up of square waves or sine waves. RPG is also capable of generating and displaying Gabor patches and raw movies/images.

RPG can be directly accessed via the shell of the Raspberry Pi, however, we expect most users will find remote accessing the Raspberry Pi via secure shell (SSH) from another computer a more user-friendly experience.

RPG is designed to be simple as possible for the user while providing flexibility. Animations are generated and saved to disk so they do not need to be created every session (Fig 2). These animations can be loaded either file-by-file or by directory, and can be triggered to display either programmatically or via 3.3V trigger (Fig 3).

**Figure 2.**
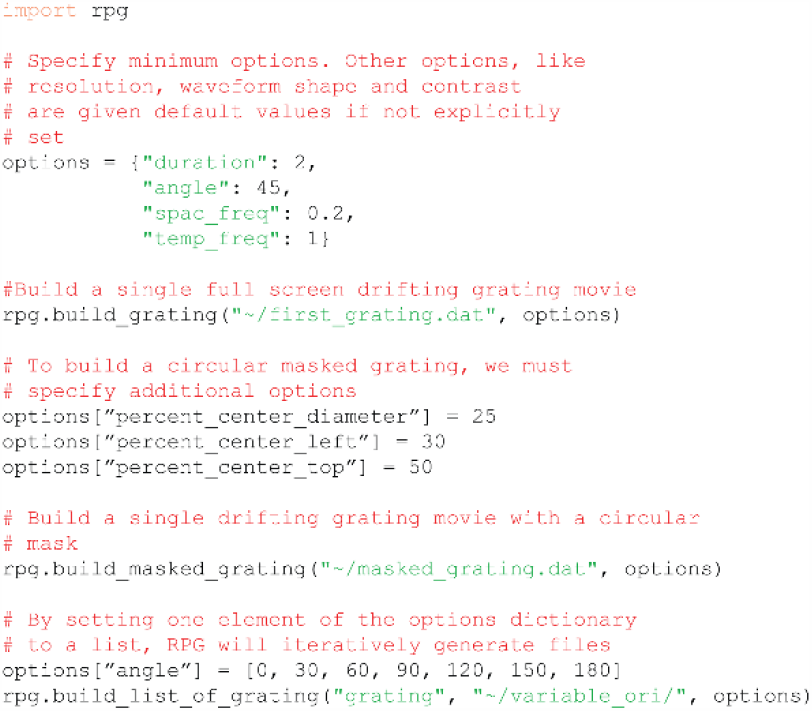
Gratings can be generated in a few lines of code, either file by file, or iteratively.

**Figure 3.**
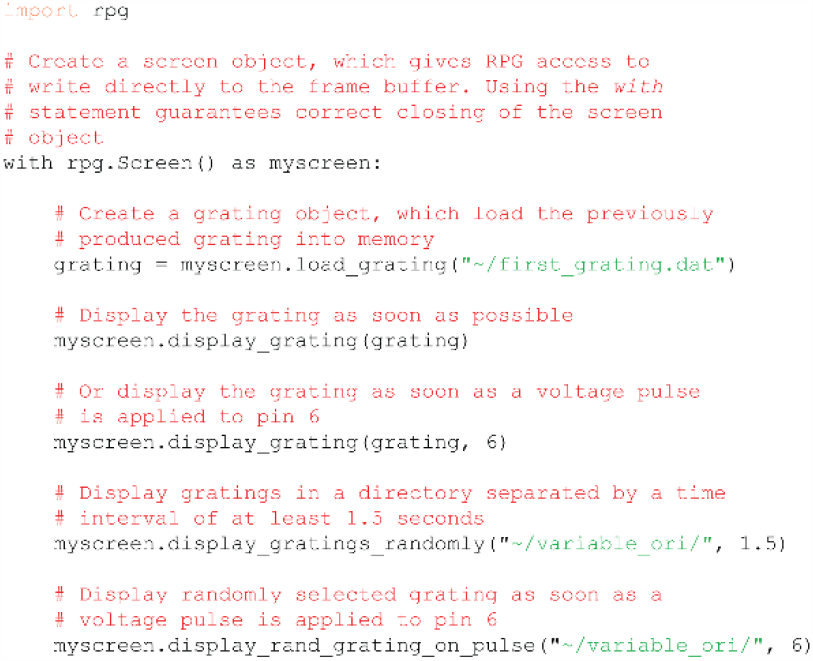
Displaying previously generated gratings can be done in as little as three lines of code. Display of gratings can be triggered programmatically or by 3.3V trigger

RPG works by directly writing to the Raspberry Pi’s “frame buffer”, the portion of memory which holds a bitmap representing all the pixel values for a complete frame of video data. This data is “double buffered”, meaning that while data is being displayed from the front buffer, RPG is writing video information to the back buffer. RPG waits until a complete frame is drawn on the monitor (v-sync) and then swaps the assignment of the front and back buffers. This behaviour prevents unwanted visual artifacts (screen tearing) and allows for accurate frame timing.

### 3.3 Expected performance

RPG delivers video stimuli with at least as dependable performance as other common packages. When measured over 3000 frames, at a requested 60 frames per second, RPG delivered frames at 59.95 frames per second, (s = 0.06 FPS). On a Windows PC (see Methods 2.2) PsychoPy delivered an equivalent stimulus at 59.98 FPS (s = 1.3 FPS), while Psychtoolbox delivered at 59.94 FPS (s = 0.6 FPS) (Fig 4C).

**Figure 4.**
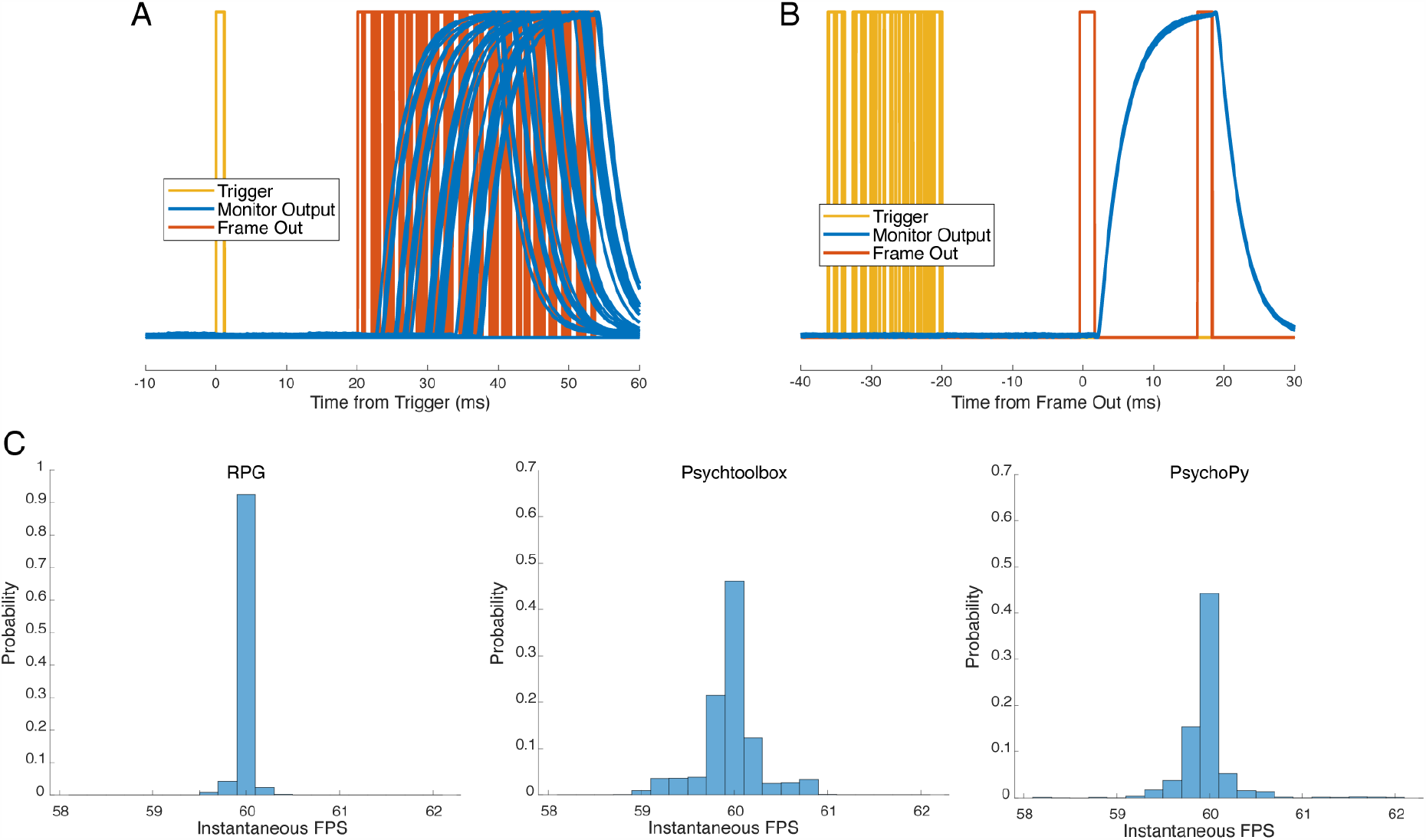
RPG’s temporal performance is at least as good as PsychoPy or Psychtoolbox. A, The delay from when RPG receives a trigger signal and when the stimulus is displayed on the monitor varies depending on where the trigger falls in the monitor refres refresh cycle. B, The response of the monitor is almost perfectly aligned to the timing of the Frame Out pulses, allowing users to reliably monitor frame timings. C, The distribution of the instantaneous frames per second (FPS), calculated as the reciprocal of the interframe time, under RPG, Psychtoolbox and PsychoPy while displaying drifting gratings.

In response to a digital trigger, RPG delivered a new frame to the monitor in as little as the duration of one monitor refresh plus 2ms, and a maximum of just under two monitor refreshes; on a 60 Hz monitor, that means as little as 19 ms, and a maximum of 33 ms (Fig. 4A). This delay depends on when the trigger signal falls relative to the timing of monitor refreshes. RPG always waits for at least one monitor refresh before updating the frame buffer, as this prevents RPG from writing to the frame buffer coincidentally with the monitor reading from it. Thus, if a trigger falls directly after a monitor refresh, RPG waits for one monitor refresh, draws to the frame buffer, which is then read and displayed on the subsequent monitor refresh. This delay can be reduced by using a monitor with higher refresh rates, e.g. using a 120 Hz monitor, where the minimum delay is 10 ms, and the maximum delay is 19 ms.

For millisecond accurate knowledge of frame timings, the “frame out” signal from RPG via physical pin 12 is essential. By directly measuring the monitor luminance, we found data appears on screen with a mean delay 2.8 ms after the frame out signal, and importantly, this delay had a standard deviation of only 15 µs (Fig. 4B). The absolute delay from the frame out going high to data appearing on the monitor is dependent on the monitor, and likely will be reduced with higher performance monitors running at higher refresh rates. This means that a user can have confidence that their stimulus was displayed on the screen a few milliseconds after the frame out pulse, but if the user performs a one-off measurement of the delay from frame out to monitor output, they can know when the stimulus was present with sub millisecond accuracy.

The built in GPIO ports of the Raspberry Pi makes it simple to trigger RPG to deliver stimuli via a TTL pulse shifted to 3.3V through voltage divider or a logic level shifter (e.g. BOB-12009, SparkFun). However, it only would only require a few additional lines of code to use the Raspberry Pi as the clock to trigger acquisition hardware/software (example provided in example/rpg_as_control.py)

### 3.4 Limitations

In order to play animations, all video data must be loaded into memory, and this leads to the primary limitation. The Raspberry Pi 3 comes with 1 GB of memory, of which approximately 750 MB is free. Each 1280 x 720 frame of 16-bit video data takes up 1.8 MB of memory, which means a maximum of approximately 400 frames can be stored in memory at one time. With no further consideration, this would result in a maximum of 7 seconds of 60 FPS animation that could be displayed without playback being paused to allow for more animations to be loaded into RAM. However, if gratings are displayed for a long duration, or at high temporal frequency, they will contain repeated frames. RPG takes advantage of this and allows animations to loop to allow for longer animations to be displayed, and hence 12 orientations of drifting gratings of 1-2 second durations of typical temporal frequencies (e.g. 2 Hz) can be stored in memory concurrently. However, natural images or movies are not compressed, and hence face the 750 MB, approximately 400 frame limit. However, the Raspberry Pi 4 is available with up to 8GB of memory. This means it can store approximately 4200 frames of 16-bit raw video, or 2800 frames of 24-bit video which is more than enough for any traditional visual experiment.

If more animations are needed than can be stored in memory, it is a simple matter to remove stored gratings from memory and load others, however, in the worst case, load times can be non-trivial. For the Raspberry Pi 3, we benchmarked gratings loading at approximately 20 MB/s from the onboard SD card or USB thumb/flash drives, meaning that it takes 2-5 times their duration to load into memory. On the other hand, using the Raspberry Pi 4 connected to an external USB 3.0 hard disk drive we bench marked loading gratings at 250 MB/s, meaning a typical grating file will load in half a second. If gratings were stored on an external solid-state drive, we expect this load time would be approximately halved. Thus, if the user wishes to play gratings rapidly in succession, while it is preferable to load all gratings in to memory before experiments begin, it is feasible to load gratings between displays using the Raspberry Pi 4, and externals USB 3.0 hard drives.

### 3.4 Expected results

Using imaging or electrophysiological software/hardware, we were able to trigger RPG to display gratings while we monitored the activity of layer 2/3 neurons in the primary visual cortex, revealing typical orientation preference behaviour (Fig 5).

**Figure 5.**
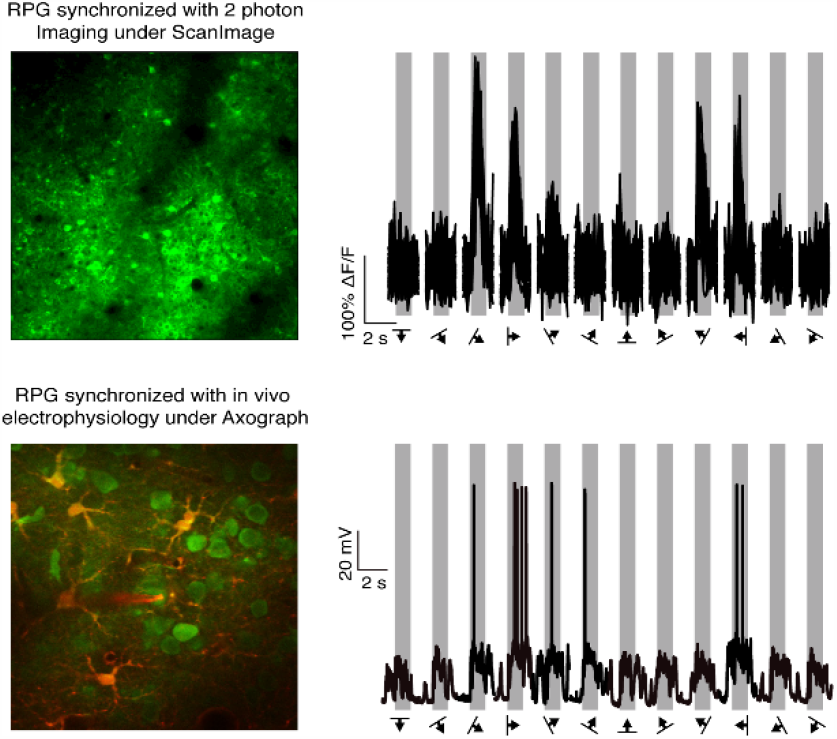
RPG is simple to synchronize with common 2-photon imaging and electrophysiology software, and produces expected cellular responses as measured with calcium imaging and in vivo whole-cell recording. Gratings were presented in a pseudo-random order.

## 4. Conclusion

RPG provides a simple, robust, flexible and high-performance solution for displaying visual stimuli. The feedback allows sub-millisecond knowledge of frame timings. When run on a Raspberry Pi 4, RPG can store large amounts of video data for rapid display, and with a USB 3.0 external hard drive, it can load data on the fly, allowing for a huge reservoir of potential stimuli. RPG is full documented and is designed to allow scientists to rapidly deploy visual stimuli with minimal difficulty.

